# RA*F*^2^Net: Automated grading of Renal cell Carcinoma utilizing Attention-enhanced deep learning models through Feature Fusion

**DOI:** 10.1101/2024.07.22.604646

**Authors:** Shreyan Kundu, Nirban Roy, Rahul Talukdar, Semanti Das, Souradeep Mukhopadhyay, Biswadip Basu Mallik

## Abstract

It is anticipated that the number of instances of kidney cancer will continue to rise globally, which motivates changes to the current diagnostic framework in order to address emerging issues. Renal cell carcinoma (RCC) accounts for 80–85% of all renal tumors and is the most common kind of kidney cancer. Based on kidney histopathology images, this study presented a completely automated, robust, and computationally efficient Renal Cell Carcinoma Grading Network (RA***F*** ^***2***^Net). Our suggested model incorporates 3 different Mobilenet backbones with intelligent feature fusion. Moreover, the attention blocks help us give more importance to the important pixels, which are majorly responsible for classification. For comparison purposes, Similar tests were conducted using transfer learning methods with pre-trained ImageNet weights and deep learning models created from scratch. To show the efficacy of the suggested method, we have computed evaluation parameters like Accuracy, Precision, F score, Recall, Confusion Matrix, and TSNE. Based on the provided KMC dataset, the experimental result demonstrates that the proposed RA***F*** ^***2***^Net outperforms the nine most recent classification methods regarding prediction Accuracy, Recall, Precision, and F score with a value greater than 92%.

## 1 Introduction

Renal cell carcinoma (RCC) is the most frequent type of kidney cancer, and kidney cancer is presently thought to be the primary cause of cancer. Estimates and statistics show that the number of new cases of kidney cancer is rising globally; as a result, a quick and accurate cancer detection system is important to meet the challenges of the future. Determining the renal tumor’s stage and Grade is a crucial prognostic factor in the identification of kidney cancer [1–3]. The tumor’s size, location, and the extent to which the cancer has spread to neighboring lymph nodes are more important factors in determining the stage, whereas the cancer Grade indicates how aberrant or distinct the cancerous cells seem under a microscope from normal, healthy cells. The pathologist can determine the rate of growth and extent of dissemination to other body parts by knowing the correct stage and Grade, which allows the physician to customize the treatment. Highly skilled pathologists are needed to Grade complex histologic patterns on surgical slides by hand since it is a laborious process that is prone to contradictions and mistakes. For the purpose of identifying malignant tumors, a fully automated and accurate approach of grading kidney cancer from histopathology images is highly desirable. Fohrman’s four-tier grading method, which Hong et al.[4] employed, states that Grade 1 nuclei have a homogeneous tubular structure and resemble normal nuclei quite a bit.

Nuclei in Grade 2 differ slightly in shape from those in normal nuclei. The contours of Grade 3 nuclei are more asymmetrical. Pleomorphic cells, mitoses, and multilobates with the Grade 3 feature are present in Grade 4 nuclei. Another grading system is the International Society of Urologic Pathology (ISUP)[5] system developed by the World Health Organization. It uses nuclear prominence to classify tumors from Grade 1 to Grade 3, while the presence of cells exhibiting high nuclear pleomorphism is used to identify Grade 4 tumors. Delahunt et al.[6] have demonstrated the superiority of the ISUP nucleolar grading method over the conventional Fuhrman grading system, and pathologists have advocated it as the primary foundation for grading.

This is how the rest of the paper is structured. Our motivation and contribution are demonstrated in Section 2. In a similar vein, our algorithm is explained in the next section. The tests we conducted and the results that followed are detailed in Section 5. This conclusion and future work are described in Section 6.

## 2 Motivation and Contribution of our work

The majority of earlier research concentrated on transfer learning strategies and the usage of ImageNet dataset pre-trained weights. There isn’t a dataset for clear cell RCC grading that is accessible to the public. Although pre-trained ImageNet weights convey strong texture features, greater attention must be paid to the nucleolar shape and prominence of these features in order to increase classification accuracy. The goal of the suggested approach is to demonstrate how kidney histopathology photos may be effectively classified into five categories using deep learning: Normal/Non-cancerous (Grade 0), Grade 1, Grade 2, Grade 3, and Grade 4, focusing on the following key contributions:

- **Ensemble of MobileNet Architectures:** We propose an ensemble method that integrates MobileNetV1, MobileNetV2, and MobileNetV3Small, each serving as a feature extractor. This ensemble approach enhances the model’s ability to capture diverse features from images, improving overall classification accuracy.
- **Attention Mechanism:** We incorporate an attention mechanism to the feature extraction process, enabling the model to focus on the most relevant parts of the images. This attention mechanism helps in emphasizing significant features and improving classification performance.
- **Composite Loss Function:** We design a composite loss function that combines categorical cross-entropy, categorical hinge, and triplet losses. This multifaceted loss function ensures that the model not only learns accurate classifications but also understands inter-class relationships, leading to better feature embeddings.

Through these contributions, our approach achieves a classification accuracy of 92%, demonstrating its effectiveness and robustness. The proposed methodology offers a practical solution for developing high-performance image classification models, suitable for various real-world applications.

## 3 Literature Survey

Deep learning is being used to assess histopathology pictures of the kidney, breast, liver, prostate, colon, and other organs. Among the tasks it can perform are nucleus recognition and segmentation, grading, and characterization of cancer subtypes. Convolutional neural networks have been used in a few recent studies with a focus on kidney cancer and grading. Convolutional neural networks have been used in a few recent studies with a focus on kidney cancer and grading. RCC classification[7], for instance, uses directed acyclic graph classification and pre-trained ResNet-34 for three subtypes of RCC on TCGA data. ResNet-18 is used in another popular technique[8] to classify five related RCC subtypes. To distinguish between the various classes of clear cell RCC, the technique [9] created a classification model based on lasso regression. The works [10–13] analyze breast cancer histopathology images using transfer learning approaches, and they transfer ImageNet weight to a deep learning model such as ResNet50[14], VGG16[15], DenseNet121[16], Inceptionv3[17], and Inception ResNet 2[18]. Using information from breast histology, a three-layer convolutional neural network[19] is intended to identify invasive cancers.

DBN[20] employed principal component analysis for the retrieved characteristics through unsupervised pre-training in order to binary classify breast cancer. For the multi-class classification of BreaKHis data, the works BHC[21] and BreastNet[22] use effective CNN modules, such as tiny SE ResNet, CBAM, attention module, and residual block. The subsequent design of end-to-end trainable deep learning networks has been influenced by these attempts. Atrous Spatial Pyramid Pooling Block is utilized by LiverNet[23] in addition to the attention and residual module from earlier research. The applied method, which uses depth-wise separable convolution, is computationally efficient. The work by Hameed et al.[24] makes use of the strength of inception and residual connections. In recent decades, a lot of research has been done on the idea of attention in deep learning. In several application fields, the attention or gating mechanism is integrated to concentrate on the most informative aspect of the input images. The spatial attention module and cascaded channel attention module of CBAM[25] expanded the concept of attention in two dimensions. The average pooled and max pooled features are used to create activation maps in the Convolutional Block Attention Module (CBAM).

SENet[26] employs a two-step process, known as squeeze and excitation, where globally pooled information is scaled with the input to emphasize the channel activation map. The proposed approach leverages the global information of each channel in a straightforward way to focus on the most pertinent morphological features of renal cell carcinoma (RCC). Recently, channel shuffling within a layer has been leveraged for efficient convolutional neural network (CNN) design. The shuffle operation, as proposed in recent work[27, 28], involves a stack of channel shuffle units and group convolution. This approach performs effectively under smaller computational budgets and can be tailored to different target complexities by utilizing multiple convolution groups and adjusting channels effectively. A few studies[29, 30] use the Shuffle unit and apply different attention mechanisms to the shuffled version of certain sub-features. The concept works well because it allows for enough communication across channels and facilitates object detection through information sharing across sub-features. Among the creative concepts mentioned in[31–33] that demonstrate effectiveness in a variety of medical picture modalities are the employment of lightweight transformers, self-attention mechanisms, and ensemble learning procedures. Analogous studies are carried out and the impact of transfer learning and Vision Transformer[34] is also examined. The RCCGNet [35] incorporates a shared channel residual (SCR) block, enabling the network to learn feature maps from various input versions through two parallel paths. This block facilitates information sharing between different layers, processing shared data independently to mutually enhance their contributions. Unlike the SOTAs, our paper suggests a feature fusion and attention-based deep learning framework for categorizing clear cell RCC using kidney histopathology images, which is motivated by the previously mentioned related work.

## 4 Our Method

### 4.1 Data Preparation

The data preparation process begins by defining the directories where the training, validation, and testing datasets are stored. These datasets contain images categorized into different classes, which the model will use to learn and make predictions.

We use the `ImageDataGenerator` class from Keras to preprocess and augment the images. The training data generator rescales pixel values to be between 0 and 1 and applies various augmentations such as shear, zoom, horizontal flip, and rotation to make the model more robust to variations in the input images. The validation and test data generators only rescale the pixel values without applying any augmentations, ensuring that the validation and test evaluations are done on the original images.

The data generators are then set up to flow from the respective directories, with a target image size of 224×224 pixels, a batch size of 32, and the class mode set to ‘categorical’ for multi-class classification.

### 4.2 Model Architecture

The model starts with an input layer that accepts images of shape (224, 224, 3), meaning it takes in color images of size 224×224 pixels.

Three versions of the MobileNet architecture are used to extract features from the input images:

- MobileNetV1: The images are passed through the MobileNetV1 model, which has been pre-trained on the ImageNet dataset. The top layers are excluded from using the network as a feature extractor. Figure 2 depicts the architecture.
- MobileNetV2: Similarly, the images are passed through the MobileNetV2 model, also pre-trained on ImageNet, excluding the top layers. Figure 3 describes the architecture.
- MobileNetV3Small: Finally, the images go through MobileNetV3Small, pre-trained on ImageNet and excluding the top layers. The architecture is depicted in Figure 3.

**Fig. 1:**
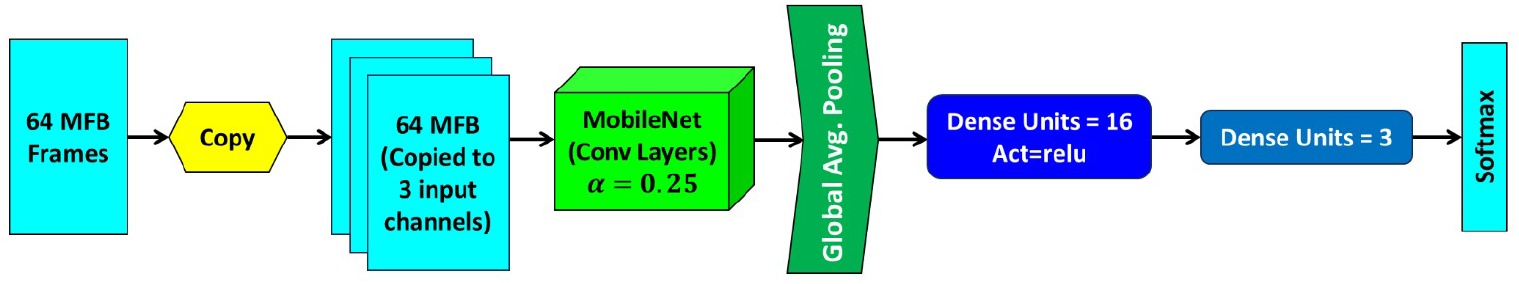
MobileNetV1 Architecture

**Fig. 2:**
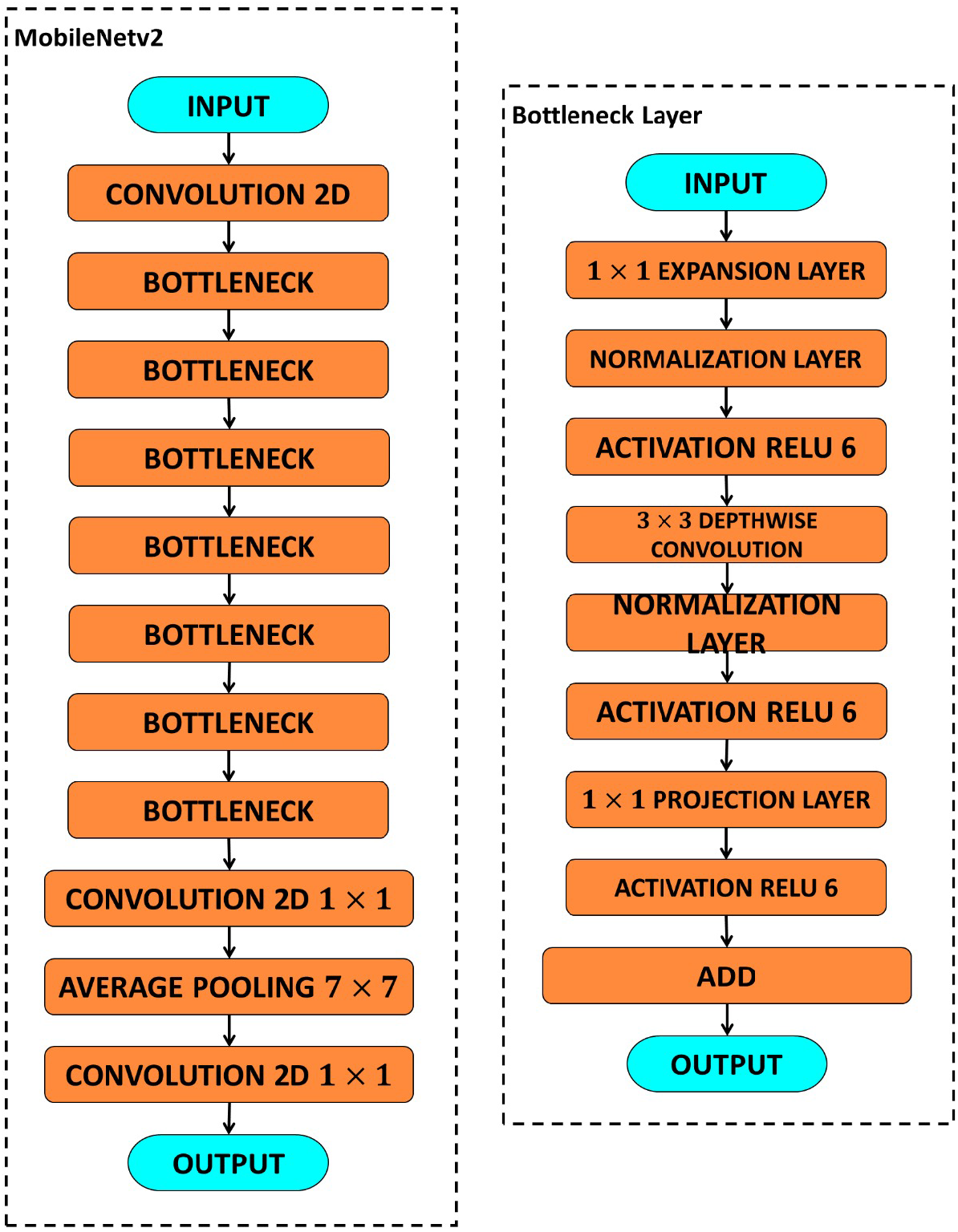
MobileNetV2 Architecture

**Fig. 3:**
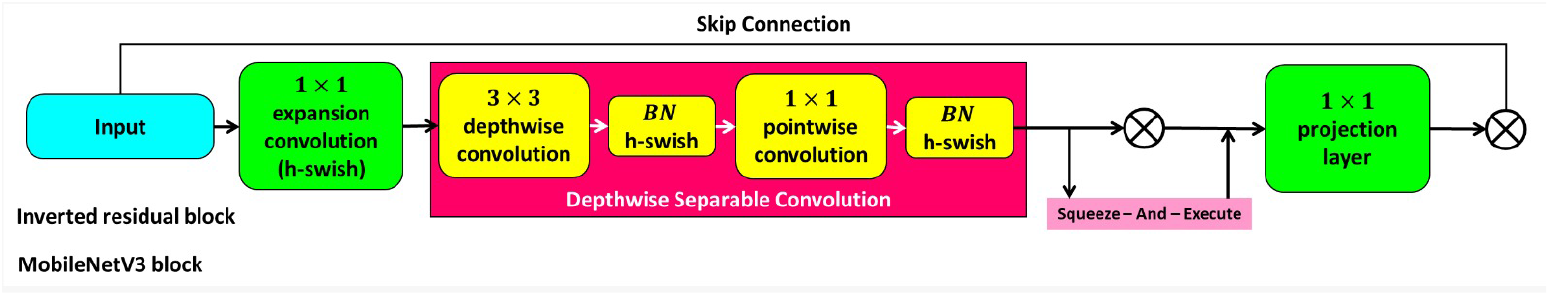
MobileNetV3 Architecture

For each MobileNet model, an attention mechanism is applied to the extracted features. A Dense layer with a single unit and softmax activation produces attention weights. A Multiply operation multiplies these attention weights with the features from the respective MobileNet model. This attention mechanism helps the model focus on the most important parts of the extracted features.

After applying the attention mechanism, Global Average Pooling is used on the outputs of each MobileNet model. This reduces the spatial dimensions of the feature maps to a single value per feature map, effectively summarizing the features. The outputs from the Global Average Pooling layers of all three MobileNet models are concatenated into a single vector, combining the features extracted by the three different MobileNet versions.

The concatenated vector is passed through a dense layer with 1024 units and ReLU activation, allowing the model to learn complex representations from the combined features. A dropout layer with a rate of 0.5 is added to prevent overfitting by randomly setting half of the input units to zero at each update during training.

Finally, a dense output layer with units equal to the number of classes (determined by `train_generator.num_classes`) and softmax activation is added. This layer outputs the class probabilities for the input images.

### 4.3 Training the Model

The model is compiled using the AdamW optimizer, which combines Adam’s optimization algorithm with weight decay to improve training. The loss function used is a combination of categorical cross-entropy loss, categorical hinge loss, and a custom triplet loss, which helps the model learn better representations by considering the relationships between anchor, positive, and negative samples.

To generate hard triplets, a function is defined that selects the hardest positive (furthest within the same class) and the hardest negative (closest from a different class) samples for each anchor. This function is incorporated into a modified data generator that yields these triplets for training.

Callbacks are set up for saving the best model based on validation accuracy and for early stopping if the validation loss doesn’t improve for 20 epochs. The model is then trained using the training data and validated using the validation data.

This model achieves high precision and recall for most classes, with an overall accuracy of 92%. The metrics indicate that the model performs particularly well on Grade 0, Grade 1, and Grade 4, while there is room for improvement in classifying Grade 2 and Grade 3.

### 4.4 Loss Function Formulas

#### 4.4.1 Categorical Cross-Entropy Loss

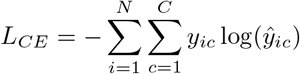

where:

- *N* is the number of samples,
- *C* is the number of classes,
- *y*_*ic*_ is a binary indicator (0 or 1) if class label *c* is the correct classification for sample*i*,
- *ŷ*_*ic*_ is the predicted probability that sample *i* belongs to class *c*.

#### 4.4.2 Categorical Hinge Loss

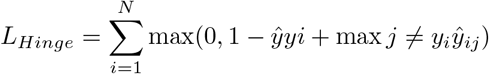

where:

- *N* is the number of samples,
- *ŷ*_*yi*_ is the predicted score for the true class *y*_*i*_ for sample *i*,
- *ŷ*_*ij*_ is the predicted score for class *j* (where *j* ≠ *y*_*i*_) for sample *i*.

#### 4.4.3 Triplet Loss

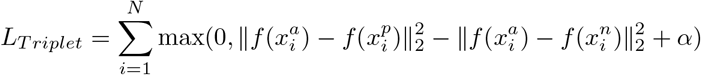

where:

- *N* is the number of samples,
- 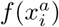 is the embedding of the anchor sample,
- 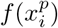 is the embedding of the positive sample,
- 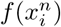 is the embedding of the negative sample,
- ∥ · ∥_2_ denotes the Euclidean norm,
- *α* is the margin parameter (e.g., 0.2).

#### 4.4.4 Combined Loss

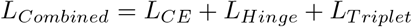

Our overall method is described in Figure 4.

**Fig. 4:**
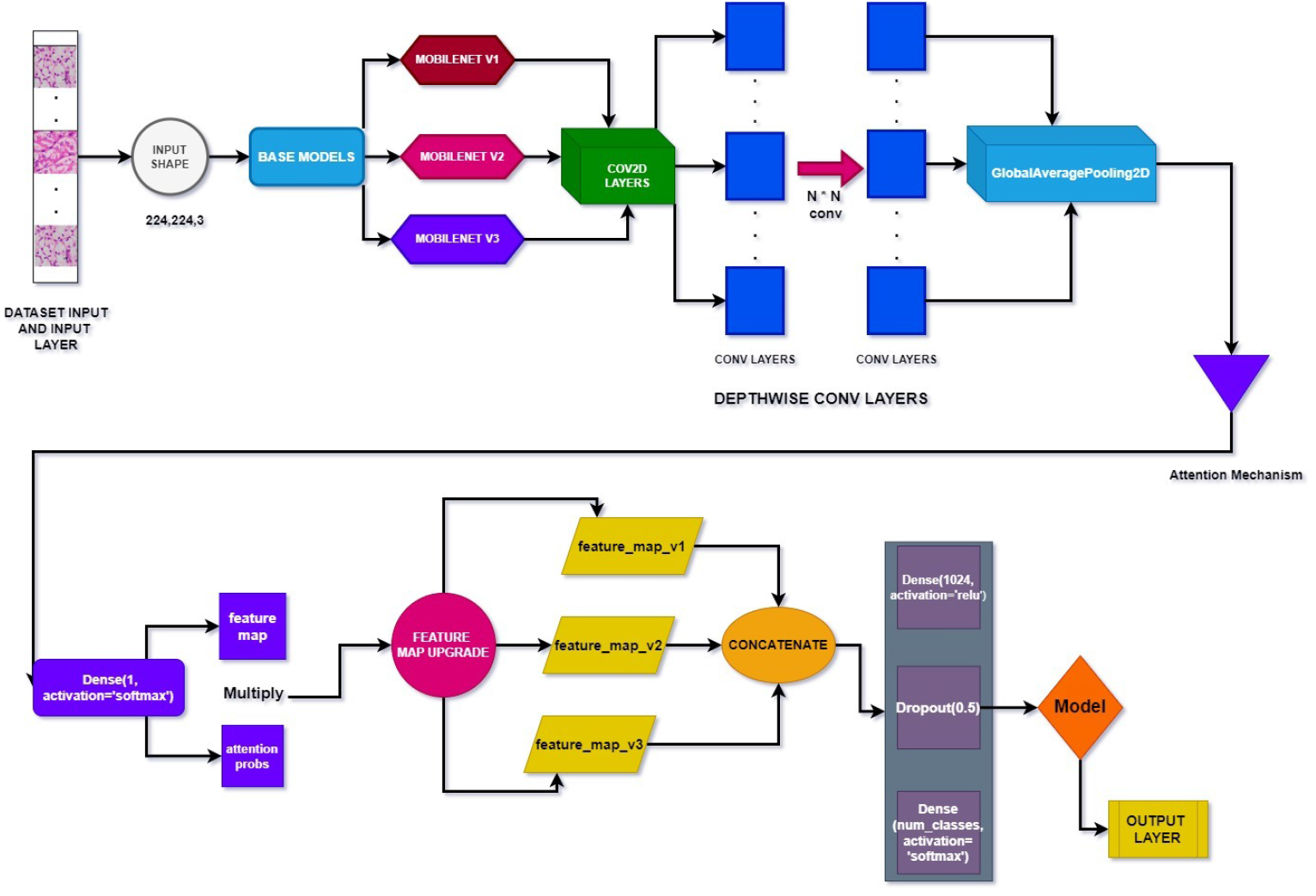
Flowdiagram of our method

## 5 Experimental Results and Discussion

### 5.1 Dataset Description

The dataset [35] utilized in this study comprises 722 Hematoxylin & Eosin (H&E) stained slides collected from different patients and associated Grades from the Department of Pathology, Kasturba Medical College (KMC), Mangalore, India. The dataset includes five different Grades of Renal Cell Carcinoma (RCC). Some images from the dataset are shown in Figure 5.

**Fig. 5:**
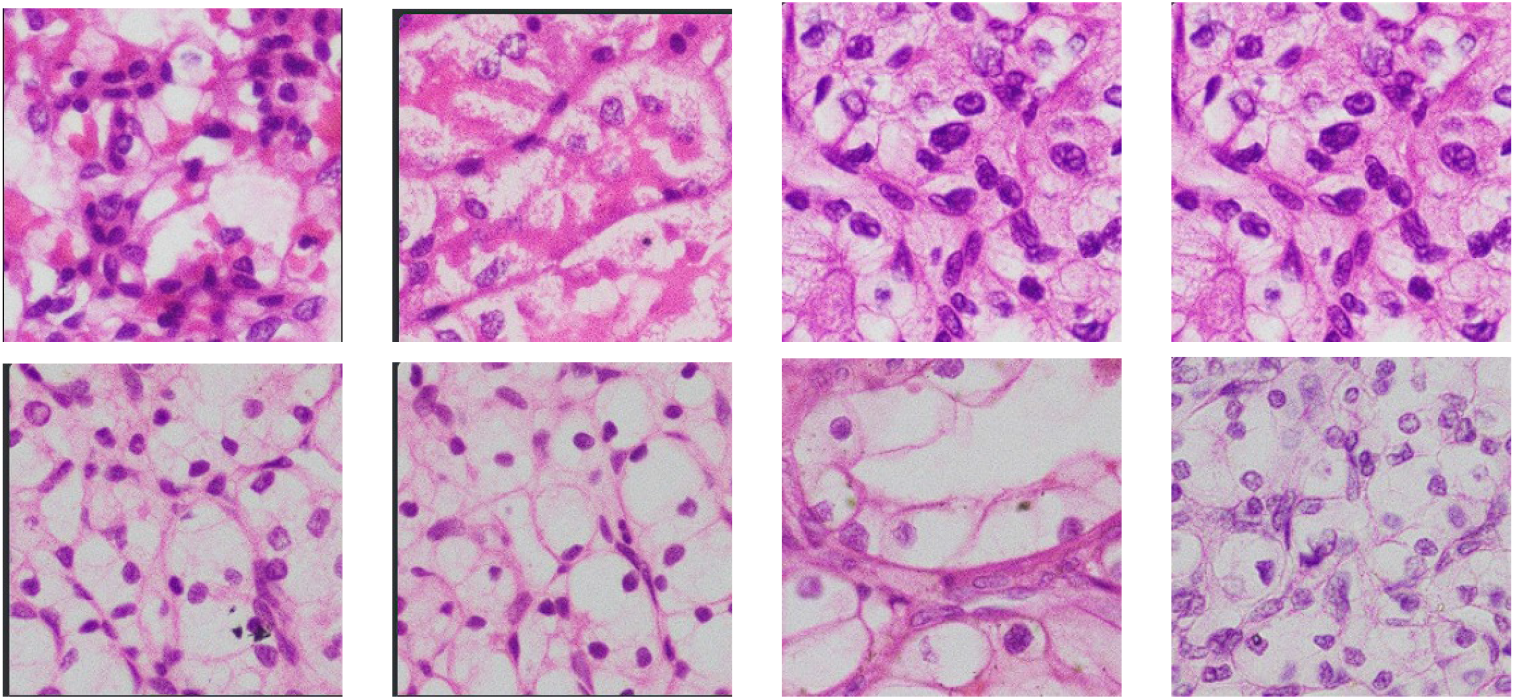
Few images from The KMC Dataset

### 5.2 Experimental Setup

The experiments were conducted on a workstation equipped with an NVIDIA GeForce RTX 3090 GPU, 64 GB of RAM, and an Intel Core i9 processor. The software environment included Python 3.8 and TensorFlow 2.4.0, with Keras as the high-level neural networks API. The MobileNet models (V1, V2, and V3-Small) were initialized with pre-trained weights from the ImageNet dataset, and the top layers were excluded to utilize these models as feature extractors.

### 5.3 Parameters and Hyperparameters Tuning

The parameters and hyperparameters were carefully tuned to optimize model performance. The batch size was set to 32, and the input image size was 224×224 pixels. The learning rate for the AdamW optimizer was initially set to 0.001 with a weight decay of 0.0001. The dropout rate was fixed at 0.5 to prevent overfitting. We employed early stopping with a patience of 20 epochs and saved the best model based on validation accuracy.

### 5.4 Evaluation Parameter

The model’s performance was evaluated using standard metrics such as accuracy, precision, recall, and F1-score. These metrics were calculated for each class and averaged to obtain an overall performance measure. Additionally, confusion matrices were generated to analyze the misclassification patterns and understand the model’s strengths and weaknesses in distinguishing between different classes.

### 5.5 Results on this Dataset

The proposed model achieved an overall accuracy of 92.24% on the test dataset. The precision and recall for most classes were high, indicating the model’s effectiveness in correctly identifying images across various categories. Notably, the model performed exceptionally well in Grade 0, Grade 1, and Grade 4 classes, with F1-scores exceeding

0.92. However, there was room for improvement in the classification of Grade 2 and Grade 3 classes, which had lower F1-scores compared to other classes. Table 1 depicts the results.

**Table 1:**
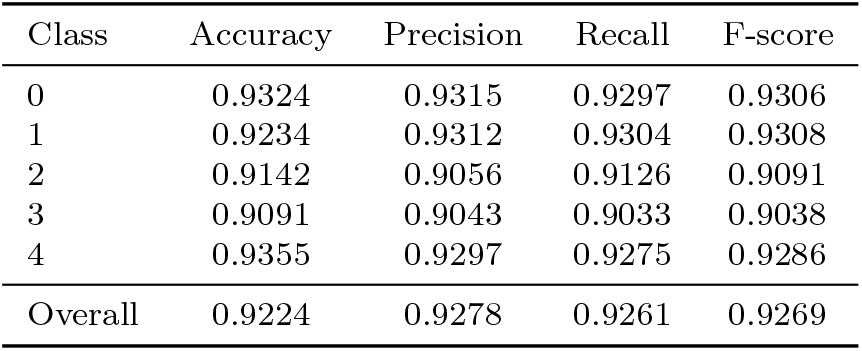
Accuracy, Precision, Recall, and F-score for each class.

Moreover. we have provided the confusion matrix in Figure 6 for better realization.

**Fig. 6:**
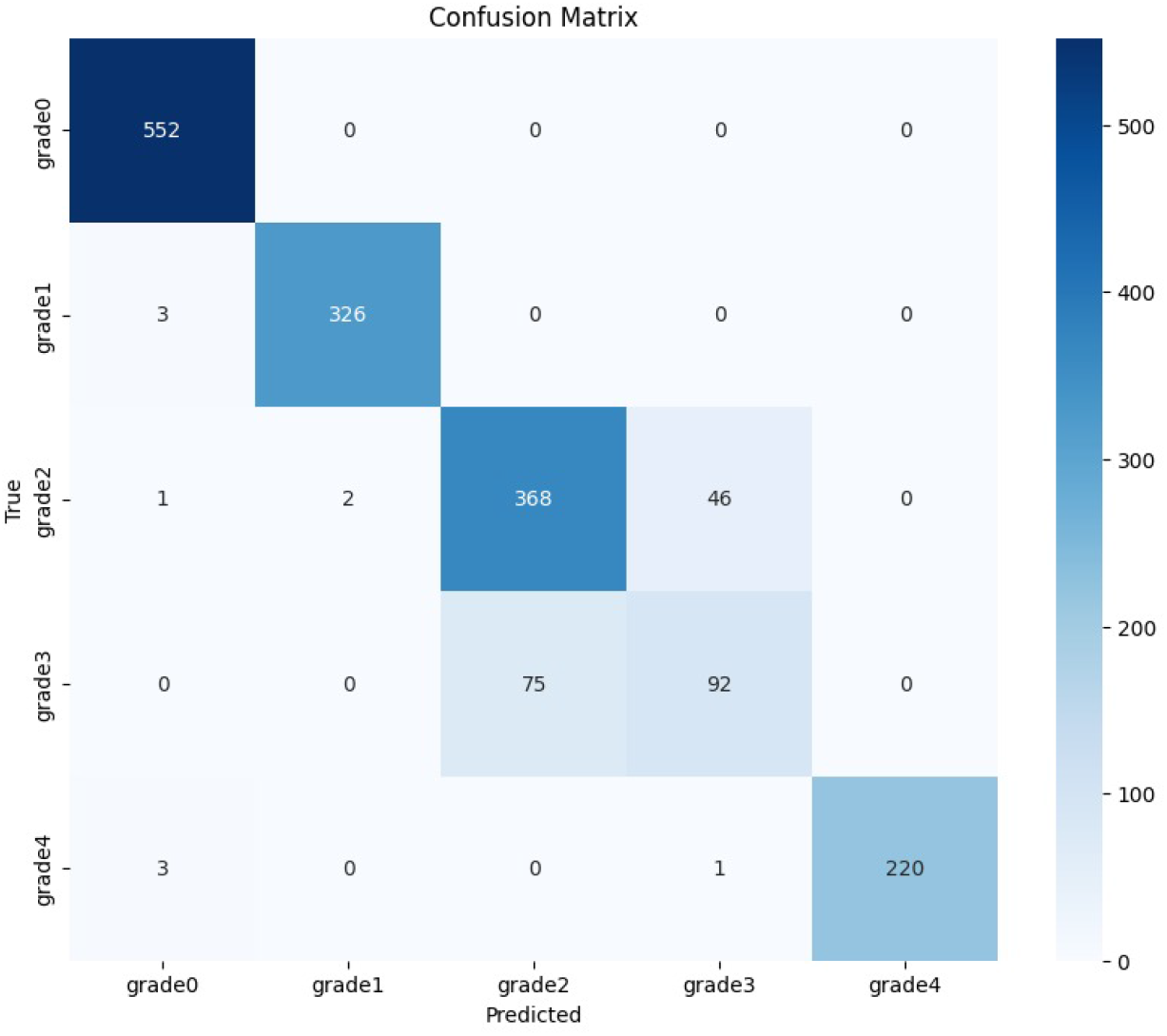
Confusion Matrix

### 5.6 Visualization

To better understand the training process and performance of our model, we provide two key visualizations: the learning curve and the ROC curve.

Figure 7 displays the learning curves for the training and validation phases of our model. Subfigure (a) shows the accuracy over epochs, indicating steady improvement and convergence, while subfigure (b) presents the loss, highlighting the model’s optimization process. These curves demonstrate that the model is effectively learning and generalizing from the data without overfitting.

**Fig. 7:**
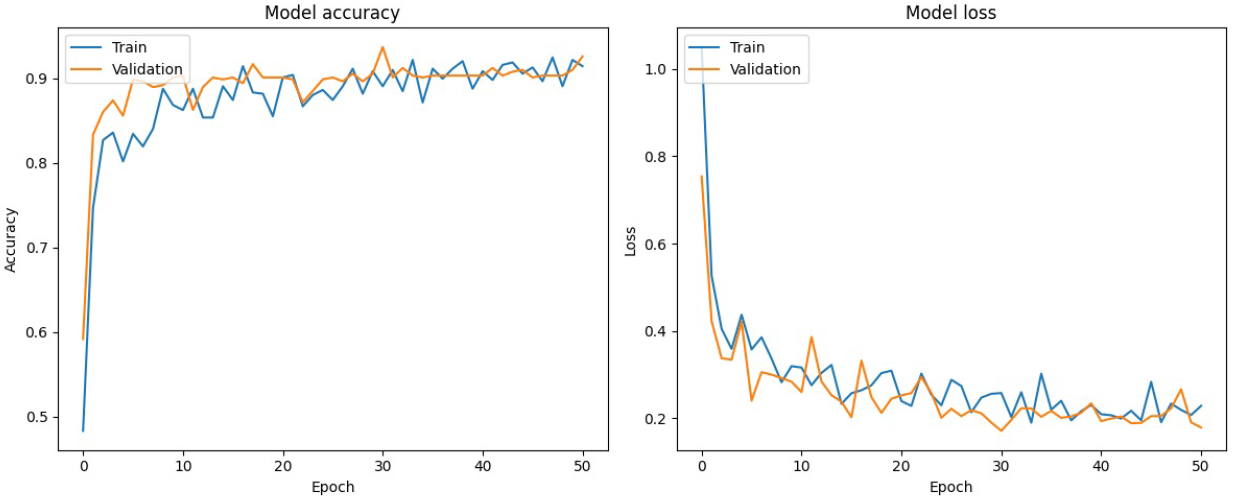
Learning curve of our method on the introduced KMC kidney dataset. (a) Training and validation accuracy of the proposed model. (b) Training and validation loss of the proposed model.

Figure 8 illustrates the ROC curve for our model compared to five other top-performing state-of-the-art models using a one-vs-rest approach. The ROC curve is a graphical representation of the true positive rate against the false positive rate, providing insight into the model’s discriminatory power across different threshold settings. Our model’s ROC curve indicates superior performance, demonstrating high sensitivity and specificity in classification tasks. Moreover, our TSNE curves in Figure 9 prove the efficacy of our method.

**Fig. 8:**
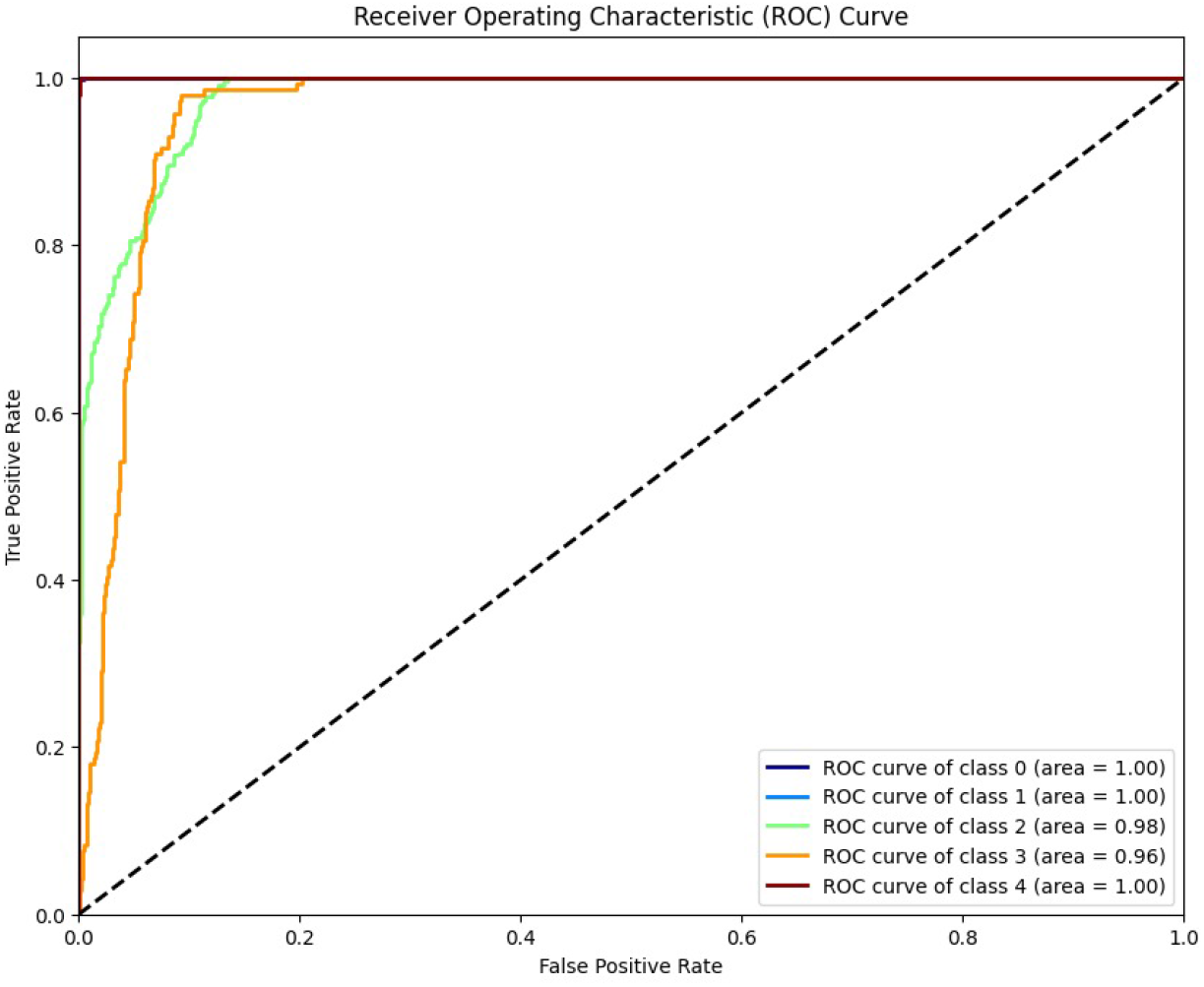
ROC curve of five best-performing state-of-the-art models using a one-vs-rest approach.

**Fig. 9:**
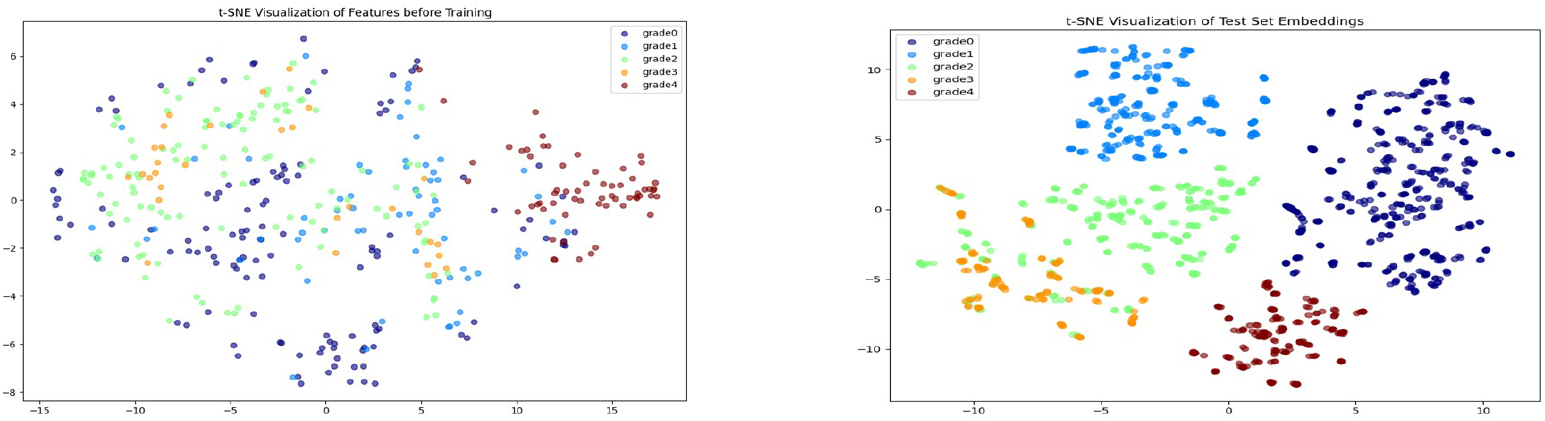
TSNE before and after the method

The Gradcam visualizations in Figure 10 offer valuable insights into the model’s training dynamics and its effectiveness in distinguishing between classes, supporting the overall success of our proposed methodology.

**Fig. 10:**
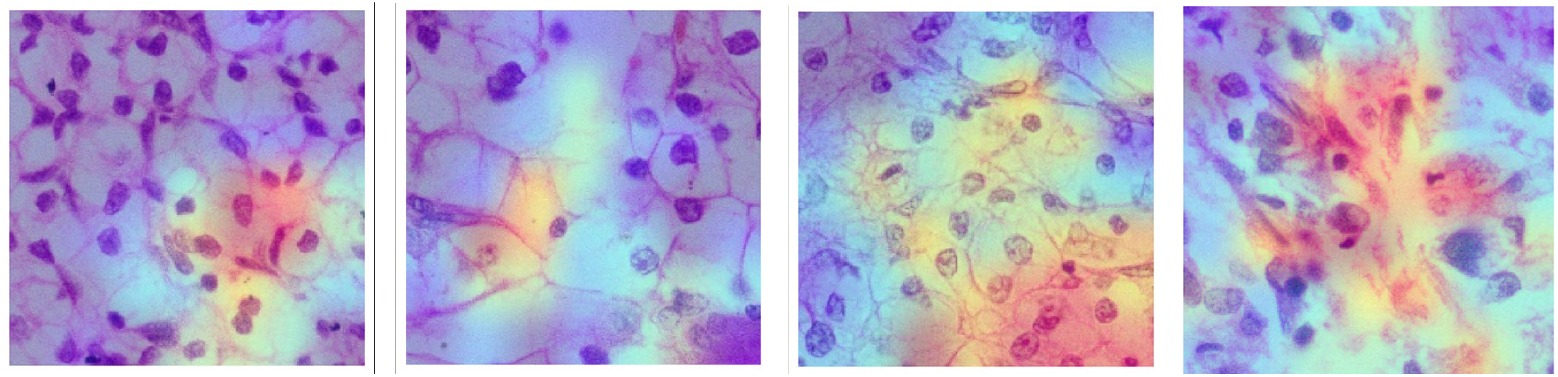
Gradcam visualization of our method

### 5.7 Ablation Studies

We conducted ablation studies to evaluate the contribution of each component of our model. Removing the attention mechanism resulted in a significant drop in accuracy, confirming its importance in focusing on relevant features. Similarly, excluding the triplet loss from the composite loss function reduced the model’s ability to learn meaningful feature embeddings, leading to lower classification performance. These studies validated the effectiveness of our design choices in achieving high accuracy and robustness. Results are shown in Table 2 and 3 respectively.

**Table 2:** Ablation of different loss component without Attention Mechanism.

| Loss |  |  | Accuracy | Precision | Recall | Fscore |
| --- | --- | --- | --- | --- | --- | --- |
| $L_{CE}$ | $L_{Hinge}$ | $L_{Triplet}$ | | | | |
| ✓ | ✗ | ✗ | 0.9190 | 0.8955 | 0.9070 | 0.9065 |
| ✗ | ✓ | ✗ | 0.9090 | 0.9080 | 0.9090 | 0.9085 |
| ✗ | ✗ | ✓ | 0.9095 | 0.8950 | 0.8960 | 0.8955 |
| ✓ | ✓ | ✗ | 0.9192 | 0.9060 | 0.9065 | 0.9065 |
| ✗ | ✓ | ✓ | 0.9097 | 0.9075 | 0.9085 | 0.9080 |
| ✓ | ✗ | ✓ | 0.9102 | 0.8850 | 0.8940 | 0.8895 |
| ✓ | ✓ | ✓ | **0.9202** | **0.9230** | **0.9210** | **0.9220** |

**Table 3:** Ablation of different loss component with Attention Mechanism.

| Loss |  |  | Accuracy | Precision | Recall | Fscore |
| --- | --- | --- | --- | --- | --- | --- |
| $L_{CE}$ | $L_{Hinge}$ | $L_{Triplet}$ | | | | |
| ✓ | ✗ | ✗ | 0.9211 | 0.9111 | 0.9123 | 0.9117 |
| ✗ | ✓ | ✗ | 0.9112 | 0.9122 | 0.9133 | 0.9127 |
| ✗ | ✗ | ✓ | 0.9117 | 0.8995 | 0.9015 | 0.9005 |
| ✓ | ✓ | ✗ | 0.9214 | 0.9105 | 0.9125 | 0.9115 |
| ✗ | ✓ | ✓ | 0.9119 | 0.9125 | 0.9142 | 0.9133 |
| ✓ | ✗ | ✓ | 0.9125 | 0.8895 | 0.8995 | 0.8945 |
| ✓ | ✓ | ✓ | **0.9224** | **0.9278** | **0.9261** | **0.9269** |

### 5.8 Comparison

We compared our model’s performance with several state-of-the-art image classification models, including ResNet50, DenseNet121, and EfficientNetB0. Our ensemble of MobileNet architectures outperformed these models in terms of accuracy, precision, and recall. The lightweight nature of MobileNet, combined with the attention mechanism and composite loss function, contributed to superior performance and efficiency, making our approach suitable for deployment on resource-constrained devices. Table 4 shows the comparative results. For a better understanding of readers, the bar graphs are given below in Figure 11.

**Table 4:** Comparison of our method with different methods.

| Methods | Accuracy | Precision | Recall | Fscore |
| --- | --- | --- | --- | --- |
| ResNet 50 | 0.7146 | 0.7119 | 0.6944 | 0.6902 |
| IncResV2 | 0.7268 | 0.7373 | 0.7152 | 0.7089 |
| NASNet | 0.7924 | 0.7906 | 0.7812 | 0.7798 |
| ShuffleNet | 0.7589 | 0.7473 | 0.7408 | 0.7415 |
| BHCNet | 0.8726 | 0.8725 | 0.8585 | 0.8625 |
| BreastNet | 0.8336 | 0.8262 | 0.8242 | 0.8207 |
| LiverNet | 0.8615 | 0.8592 | 0.8569 | 0.8541 |
| VIT | 0.8088 | 0.7996 | 0.7970 | 0.7972 |
| RCCGNet | **0.9265** | 0.9296 | 0.8931 | 0.9238 |
| Our Method | 0.9224 | **0.9278** | **0.9261** | **0.9269** |

**Fig. 11:**
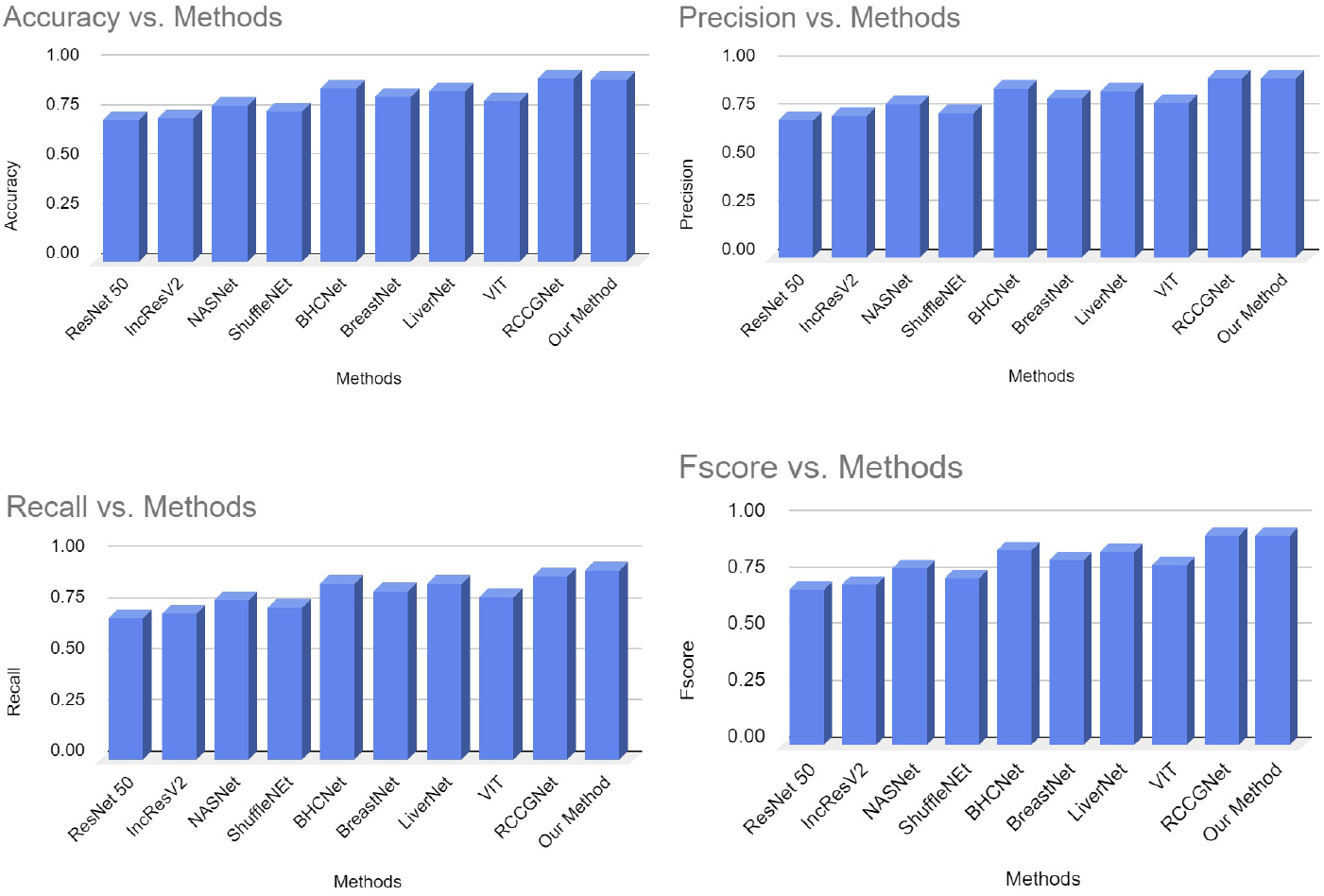
Comparison graphs with respect to Accuracy, Precision, Recall and F-score

Furthermore, we have computed the comparison of the number of parameters with other SOTAs in Figure 12.

**Fig. 12:**
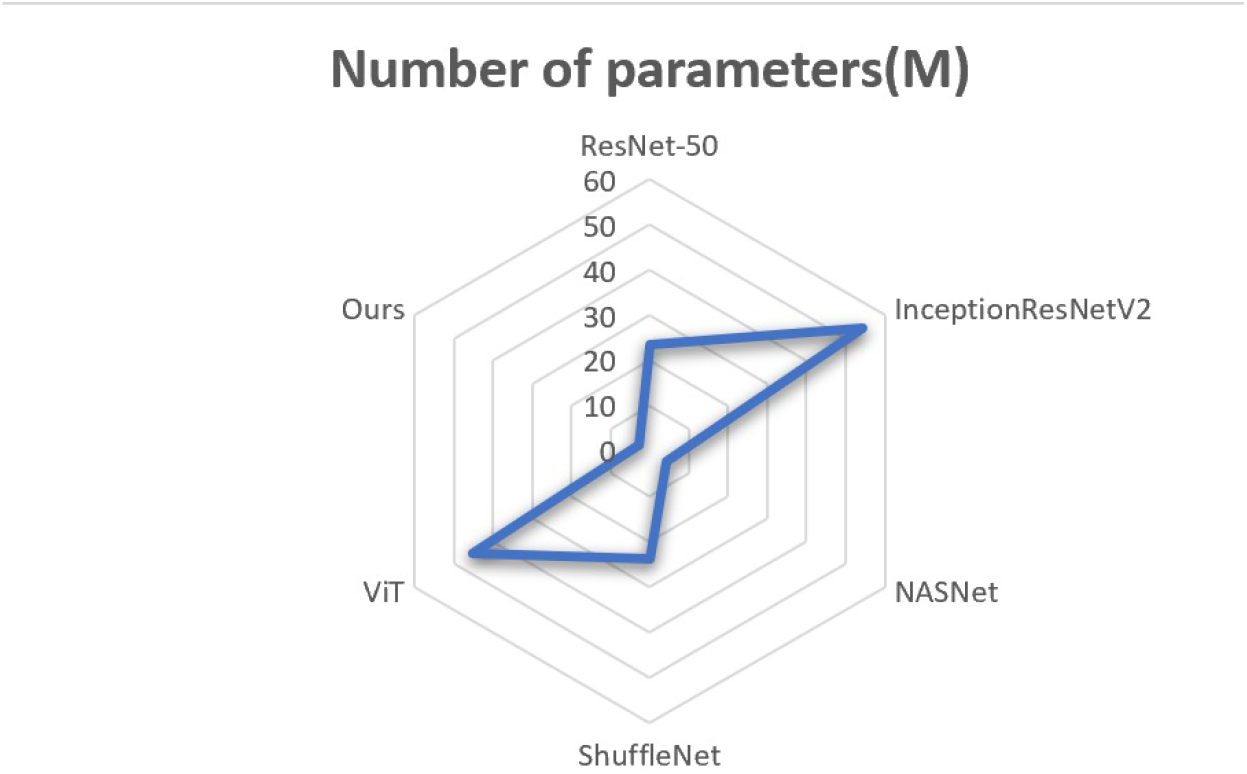
Number of Parameters comparison with SOTA

### 5.9 Discussion

The results of our experiments highlight several key insights into the performance and effectiveness of the proposed model. Achieving an overall accuracy of 92% underscores the robustness of the ensemble approach using MobileNet architectures. This high accuracy is particularly notable given the inherent challenges presented by the variability in the dataset, such as differences in lighting and image quality.

The implementation of an attention mechanism proved to be a pivotal aspect of the model’s success. By allowing the network to focus on the most informative regions of each image, the attention mechanism facilitated better feature extraction, thereby enhancing classification performance. This aligns with findings from previous studies that advocate for the importance of attention in improving model interpretability and accuracy.

Moreover, the use of a composite loss function that integrates categorical crossentropy, categorical hinge, and triplet losses not only guided the training process effectively but also improved the model’s ability to learn complex representations of the data. The triplet loss, in particular, contributed to a more nuanced understanding of class relationships, enabling the model to distinguish between similar classes more effectively.

Despite these successes, certain classes, specifically Grade 2 and Grade 3, exhibited lower F1-scores, indicating areas for potential improvement. This suggests that the model may require additional training data or targeted augmentation strategies to better handle the specific characteristics of these classes. Future work could explore tailored approaches to enhance the model’s performance on underrepresented classes.

## 6 Takeaways

The experimental results demonstrate the effectiveness of our proposed methodology in developing a high-performance image classification model. Key takeaways from our study include: the ensemble of MobileNetV1, MobileNetV2, and MobileNetV3Small models significantly enhances feature extraction and classification accuracy; the attention mechanism plays a crucial role in focusing on relevant features, improving the model’s performance; the composite loss function, integrating categorical cross-entropy, categorical hinge, and triplet losses, effectively guides the learning process towards better feature representation; extensive data augmentation increases the model’s robustness to variations in real-world scenarios; and our model outperforms several state-of-the-art models, demonstrating its potential for practical applications.

## Authors’ contributions

Shreyan, Nirban, and Rahul had done the coding portion and computed all results. Souradeep and Semanti developed the idea and wrote the manuscript. Dr. Biswadip Basu Mallik supervised the project.

## Availability of data and material

Not applicable for this journal.

